# A cross-environment comparison of nontuberculous mycobacterial diversity

**DOI:** 10.1101/2025.08.04.668438

**Authors:** Matthew J. Gebert, Ettie M. Lipner, Jordan M. Galletta, Jessica B. Henley, Michael Hoffert, Melissa L. Riskin, D. Rebecca Prevots, Noah Fierer

**Affiliations:** Department of Ecology and Evolutionary Biology, University of Colorado, Boulder, CO, USA; Cooperative Institute for Research in Environmental Sciences, University of Colorado, Boulder, CO, USA; Epidemiology and Population Studies Section, Division of Intramural Research, National Institute of Allergy and Infectious Diseases, National Institutes of Health, Rockville, MD, USA; U.S. Geological Survey, Hydrologic Networks Branch, Lawrenceville, NJ, USA

**Keywords:** mycobacteria, NTM, nontuberculous mycobacteria, *Mycobacterium*, premise plumbing

## Abstract

Nontuberculous mycobacteria (NTM) are a group of environmental bacteria that encompass nearly 200 described species, some of which can cause chronic pulmonary infection in humans. What makes these infections unique is that they are environmentally acquired, yet there remains a limited understanding of how different environments contribute to potential pathogen exposure. Here, we use new and existing marker gene datasets to compare the amounts and types of NTM across three environments known to harbor mycobacteria, surface waters, soil, and household plumbing biofilms, to better understand potential pathogen occurrence in each environment. We used 16S rRNA gene sequencing, in tandem with mycobacterial-specific marker gene sequencing to characterize variation in the relative abundances of the genus *Mycobacterium* and specific mycobacterial taxa across the three environments, with a focus on a clinically significant NTM. We found that household plumbing biofilms contained both the highest relative abundance of the genus *Mycobacterium* (on average, 13.7% of bacteria were members of the genus), as well as the highest occurrence of clinically relevant species detected (*Mycobacterium avium*, *Mycobacterium abscessus*), compared to surface waters and soil. Although mycobacteria are ubiquitous across many different environments, mycobacterial diversity is highly variable between environments with clinically relevant species largely restricted to household plumbing biofilms, information that is critical for understanding the ecology and epidemiology of NTM disease.

**Importance:** Nontuberculous mycobacteria, or NTM, are a diverse group of bacteria within the genus *Mycobacterium* that are common in many environments. While most members of the genus pose little threat to human health, a handful of species, namely the *Mycobacterium avium* complex, *M. abscessus*, and *M. mucogenicum*, can cause severe and prolonged lung infections. These environmentally acquired infections are on the rise in the United States and around the world, yet we still don’t have a good understanding of which environment types pose the greatest risk of infection to susceptible populations. Our study used cultivation independent approaches to identify the specific NTM taxa found in over 1,000 samples from three potentially important environmental reservoirs - surface waters, soils, and household plumbing systems, to determine which of these environments are most likely to harbor NTM of clinical significance. Our results highlight the high degree of variability in the types of NTM taxa detected in different environments (including extensive novel diversity within the genus) and show that household plumbing biofilms are likely the most important reservoir and subsequent route of transmission for clinically significant NTM.

## Introduction

Nontuberculous mycobacteria (NTM) are a group of bacteria commonly found in a wide range of environments. They are defined by their unique cell wall structure, consisting of long-chain fatty acids known as mycolic acids, that allow for their prolonged survival during periods of desiccation and nutrient limitation, as well as resistance to common disinfectants and antibiotics (Brennan and Nikaido 1995). Although most members of the genus are of limited clinical significance, some species (e.g. *Mycobacterium avium* complex, *Mycobacterium abscessus*, *Mycobacterium kansasii*) can cause pulmonary infection in humans, especially in individuals with underlying structural lung damage (Lake et al. 2016). NTM disease is of growing concern worldwide, with case numbers increasing annually across the United States and around the world (Winthrop et al. 2020; Dahl et al. 2022; Bents et al. 2024).

NTM disease is widely considered to be environmentally acquired, with many environments implicated in the exposure and transmission of pathogenic NTM (Falkinham 2016; Gebert et al. 2018; Gross et al. 2021). While NTM have been detected in a wide range of environments, including soil, household plumbing, hot tubs and pools, and natural water bodies (Centers for Disease Control and Prevention, Atlanta, GA, United States 2024), few studies have comprehensively characterized NTM communities across a range of environments or quantified the occurrence of clinically relevant NTM taxa within different types of environments. This knowledge gap persists, in part, because of the widespread reliance on cultivation-dependent approaches for assessing NTM diversity in environmental samples, and the biases and limitations associated with such approaches (Modra et al. 2023; Ulmann, Kracalikova, and Dziedzinska 2015; von Reyn et al. 1993; Bland et al. 2005; Mercaldo et al. 2023). Many environmental NTM remain undescribed or poorly characterized (Walsh et al. 2019), and differentiating clinically relevant NTM from those that are not clinically relevant requires time-consuming genotyping of isolates (Baldwin et al. 2019; Sarro et al. 2018). Likewise, previous work characterizing the presence of clinically relevant NTM in different environments has often focused on relatively small numbers of representative samples (Jacobs et al. 2009; Kopecky et al. 2011; Roguet et al. 2016), making it challenging to reach broader conclusions about the potential importance of different environments as sources of NTM infections. In short, much of our knowledge on which environments most likely harbor clinically relevant NTM is derived from limited or anecdotal evidence.

Previous studies have used cultivation-independent approaches to survey NTM diversity in two environments often implicated as reservoirs of clinically relevant NTM: premise plumbing and surface soils. More specifically, Gebert et al. analyzed showerhead biofilms collected from 638 homes across the US (Gebert et al. 2018) and Walsh et al. analyzed 143 soils from across the globe (Walsh et al. 2019), using a combination of 16S rRNA and *hsp65* gene sequencing of DNA extracted directly from the environmental samples to characterize NTM diversity and composition. Both survey efforts found that the NTM communities found in these environments are diverse and highly variable in composition with clinically relevant NTM frequently detected in building premise plumbing. Surface waters, including lakes, streams, and reservoirs, are also implicated as sources of NTM exposures, and are known to harbor a broad diversity of NTM, including clinically relevant taxa.

(Delghandi et al. 2020; Roguet et al. 2016; George et al. 1980; Makovcova et al. 2014). However, it remains unclear how NTM communities vary across surface waters and, whether surface waters represent an important source of exposures to clinically relevant NTM. More generally, it remains unclear how NTM diversity differs across these environments and which environments represent the most important potential sources of NTM exposures.

For this study, we used high throughput marker gene sequencing, targeting both the 16S rRNA gene and the mycobacterial-specific 65-kD heat shock protein (*hsp65*) gene to characterize the ubiquity and types of NTM detected across 373 surface water samples collected from 102 sites across the US. We then integrated the surface water datasets with previously published datasets from 143 soils (Walsh et al. 2019) and 638 household plumbing biofilm samples (Gebert et al. 2018) to directly compare the diversity and ubiquity of NTM found across these three major environment types. The comprehensive survey of NTM diversity in surface waters conducted here, when coupled with the soil and premise plumbing datasets, made it possible to quantitatively compare NTM communities across environments and identify which environments are most likely to harbor clinically relevant NTM.

## Methods

### Surface water samples

A total of 373 surface water samples were collected from surface water sites across the United States that are monitored by the US Geological Survey (USGS) as part of the National Water Quality Network (NWQN) (Riskin and Lee 2021). These samples were collected from 102 unique surface water collection sites across 28 states (for site map, see Figure 1). At 73 of the sites, multiple samples were collected over time starting in July 2022 and concluding in November 2022. The collection sites spanned 8 different surface water types, with the majority coming from streams, but also including large inland and coastal rivers, lakes and reservoirs, and urban and agricultural sites (Supplementary Table 1). For the 16S rRNA gene sequencing analysis, all samples (N = 373) were included in downstream analysis.

**Figure 1:**
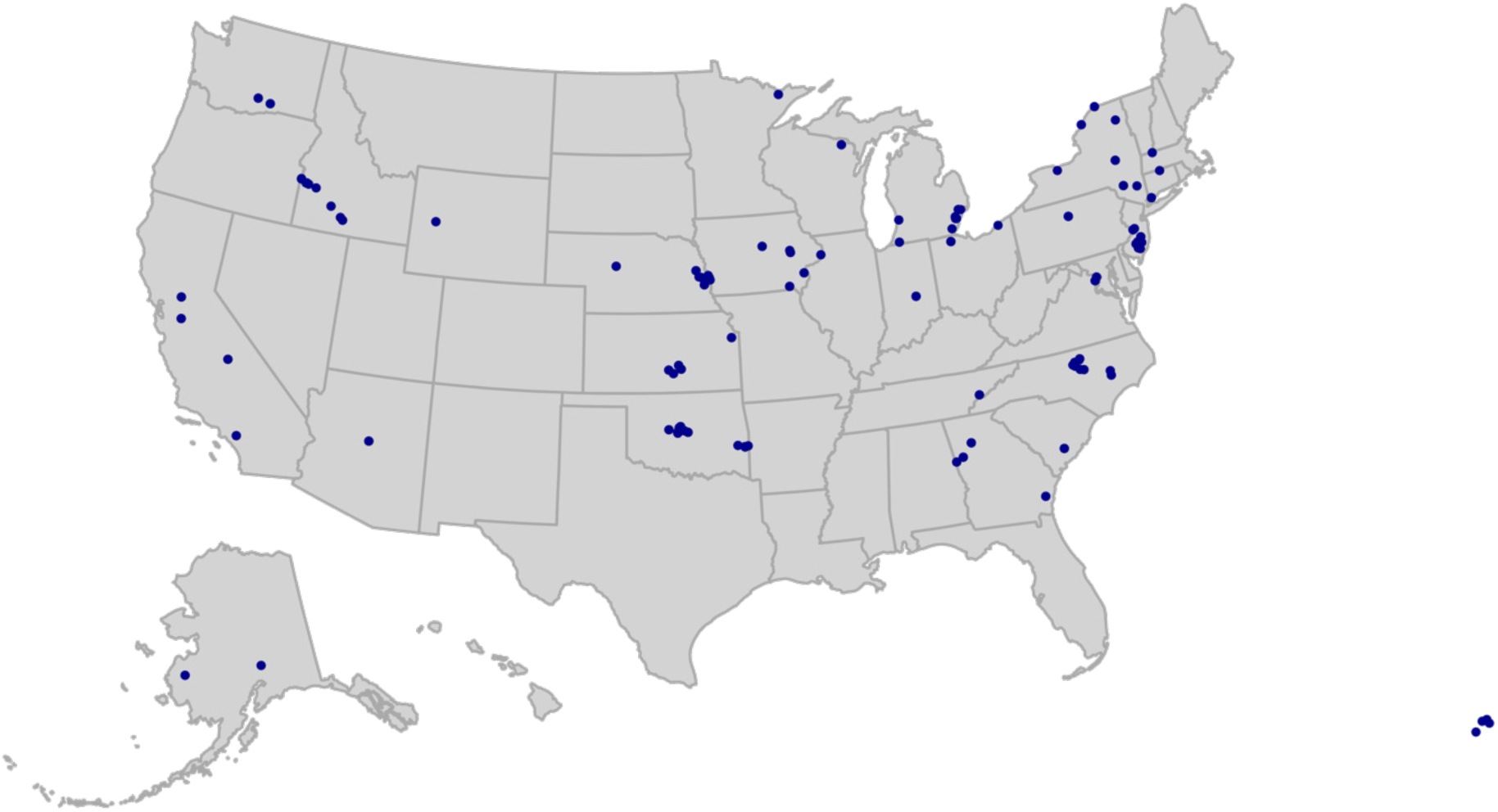
Map showing the distribution of surface water sample collection sites, colored in purple, across the United States (N = 102), as part of the National Water Quality Network (USGS). For information on collection locations for the soil and household plumbing biofilm samples, see Walsh et al. (2019) and Gebert et al. (2018).

Methods of sample collection used by the NWQN conform to the USGS National Field Manual for the Collection of Water-Quality Data (US Geological Survey 2006). To the greatest extent possible, isokinetic, depth-integrated sampling techniques that provide samples representative of stream conditions are used. The collection of isokinetic, depth-integrated samples is done using either an equal-width-increment (EWI) or equal-discharge-increment (EDI) sampling method that yields a composite sample that represents the streamflow-weighted concentrations of the stream cross section being sampled. For each sample collection, a total of 0.2L - 10L of water was filtered immediately through 0.45 µm GWV high-capacity groundwater sampling capsule filters (Cytiva Life Sciences) by USGS personnel. Collection filters were then stored at -20°C at individual sites until the final date of collection (November 2022). All capsule filters were shipped overnight on ice to the University of Colorado Boulder, where they were stored at -20°C until DNA extraction.

Metadata associated with all samples were downloaded from the Water Quality Portal (https://www.waterqualitydata.us/) and included information on water temperature (°C), water pH, dissolved oxygen (% saturation), total carbon (mg/L), total nitrogen (mg/L), sodium adsorption ratio (SAR), and turbidity at the time of sampling (Supplementary Table 3).

### Household plumbing biofilm samples

We re-analyzed a previously published dataset from a large-scale survey of showerhead biofilms, treating the showerhead biofilms as a proxy for household plumbing biofilms (Gebert et al. 2018). For this analysis, we focused only on those biofilm swab samples collected from households in the United States (N = 638, 49 of 50 states in the US represented). For specific site information and sampling details, see Gebert et al. (2018).

### Global soil samples

We also re-analyzed a previously published soil dataset from Walsh et al. (2019), which focused on a subset of 143 soils from collection sites across 6 continents, originally collected for the study described in Delgado-Baquerizo et al. (2018). Composite samples were collected from each site to 7.5 cm depth with all samples stored at -20°C immediately after collection. For additional soil and site information, see Delgado-Baquerizo et al. (2018) and Walsh et al. (2019).

### DNA extraction

DNA was extracted from the showerhead biofilm and soil samples using the DNEasy PowerSoil kit (Qiagen, Germantown, MD, USA), see Gebert et al. (2018) and Walsh et al. (2019) for further details. To extract DNA from the surface water samples, filters were first thawed at room temperature for ∼1 hour prior to downstream processing. Autoclaved 7/16” brass caps were attached to the bottom of each filter capsule prior to thawing. Immediately before shaking, 5mL of 5% Triton was added to each filter capsule, and the filter was capped on the top end, sealing the capsule. Capsule filters were shaken at 20 Hz for 7.5 min, then inverted and shaken for another 7.5 min. We then used an air-filled 50mL syringe to force liquid out of each capsule in the direction of flow, collecting 2-5 mL of eluent per capsule into a DNA-free 15 mL conical tube, with the eluents frozen at -20°C immediately after collection. After thawing the eluents at room temperature, we extracted DNA using the DNeasy PowerWater kit (Qiagen, Germantown, MD, USA), following the manufacturer’s recommended protocol. In each round of extraction, we included an extraction blank to check for contamination.

### 16S rRNA Gene Sequencing

We amplified and sequenced a portion of the 16S rRNA gene using bacterial and archaeal primers to determine the relative abundance of the genus *Mycobacterium* in each sample. For a detailed description of library preparation and sequencing for the soil and household plumbing biofilm samples, see Walsh et al. (2019), and Gebert et al. (2018). To characterize the bacterial community of the extracted surface water, we used the same barcoded 515fBC/806rB primers (Caporaso et al., 2012) as aforementioned, which target the V4 region of the 16S rRNA gene. Duplicate PCR reactions were performed as follows: 12.5μl of Colorless Platinum II Hot Start Master Mix (ThermoFisher, Waltham, MA, USA), 10μl of Sigma water (Sigma-Aldrich, Burlington, MA, USA), 1μl of primers and 1μl of gDNA. Thermocycler conditions were 94°C for 2 minutes, 35 cycles at 94°C for 15 seconds, 60°C for 15 seconds, 68°C for 1 minute, then 72°C for 10 minutes. Amplicon normalization and clean-up was done using the Invitrogen SequalPrep Normalization Plate kit (ThermoFisher, Waltham, MA, USA) with 5μl of amplicons from each sample pooled. The composited library was then quantified using the Invitrogen Qubit dsDNA HS Assay kit (ThermoFisher, Waltham, MA, USA) and ABsolute qPCR Mix, SYBR Green, no ROX (ThermoFisher, Waltham, MA, USA). The library was run on the Illumina MiSeq using a v2-300 cycle kit (Illumina, San Diego, CA, USA) at the Center for Microbial Exploration at the University of Colorado Boulder.

### Hsp65 gene sequencing

As 16S rRNA gene sequence data are not generally useful for identifying particular mycobacterial taxa (Kim and Shin 2018), we also sequenced the 65-kD heat shock protein (*hsp65*) gene using mycobacterial-specific primers Tb11-Tb12 primers (Telenti et al. 1993). For a detailed description of library preparation and sequencing for soil and household plumbing biofilm samples, see Walsh et al. (2019), and Gebert et al. (2018). For the surface water DNA extracts, the same protocol described in Gebert et al. (2018) was followed except we used Platinum II Hot Start Master Mix (ThermoFisher, Waltham, MA, USA) for PCRs and amplicons were cleaned by adding 8.85 μL DNA-free water and 0.251 μL of ExoSAP (New England Biolabs, Ipswich, MA) to a 20 μL aliquot of each amplicon followed by incubation at 37°C for 30 minutes followed by 95°C for 5 minutes.

A second PCR was done to attach 12-bp universal reverse barcodes for sample multiplexing. Each reaction consisted of 12.5 μl of Platinum Hot Start Master Mix (ThermoFisher, Waltham, MA, USA), 8.5 μL of Sigma water (Sigma-Aldrich, Burlington, MA, USA), 2 μl of combined Illumina universal barcoded primers stock at 10μM each (Illumina, San Diego, CA, USA), and 2 μL of cleaned amplicon from the first reaction, with thermocycler conditions as follows: 95°C for 3 minutes, then 8 cycles of 95°C for 30 seconds, 55°C for 30 seconds, 72°C for 30 seconds, then 72°C for 2 minutes and held at 4°C. We followed the same protocol as above for normalization and quantification of the library. Sequencing was done using a v3-600 cycle kit (Illumina, San Diego, CA, USA) on the Illumina MiSeq at the Center for Microbial Exploration at the University of Colorado Boulder.

### 16S rRNA Gene Sequence Analyses

Raw reads from the surface water samples were demultiplexed using idemp (https://github.com/yhwu/idemp) and primers were trimmed from reads using cutadapt (version 1.8.1)(Martin 2011). Data was processed using the DADA2 pipeline (version 1.22.0)(Callahan et al. 2016). Reads were filtered based on the quality metrics (trunclen = (150, 140), maxEE = (2,2), truncQ=2, maxN=0). After the removal of chimeras, taxonomy was assigned against the SILVA nr v.138.1 database (Quast et al. 2013). This resulted in 376 samples being used for downstream analysis. The 16S rRNA marker gene datasets from the two previously published studies (household plumbing biofilms and soil) were generated using an identical procedure (Gebert et al, 2018, Walsh et al, 2019). After processing and quality filtering, a total of 638 household plumbing biofilm samples and 143 soil samples were included in this study. Combined with the 373 surface water samples that met our threshold for inclusion (minimum 2,000 archaeal and bacterial 16S rRNA gene reads per sample), the combined 16S rRNA gene dataset used for downstream analyses included a total of 1,154 samples.

### Hsp65 Sequence Analyses

The *hsp65* marker gene sequence data from each of the three datasets were combined and processed together using a single pipeline to maintain consistency and permit direct comparison of results across the three unique environments. First, each *hsp65* dataset was demultiplexed independently using idemp (https://github.com/yhwu/idemp). Primers were trimmed from reads using cutadapt (version 1.8.1) (Martin 2011). Data was then processed using the DADA2 pipeline (version 1.22.0) (Callahan et al. 2016). Each dataset was filtered independently based on the quality metrics for each respective MiSeq run. Error rates were then determined, and sequence variants were inferred (pool = TRUE), resulting in three amplicon sequence variant (ASV) tables, one for each environment. Sequence tables from each of the three independently processed datasets (household plumbing biofilms, global soils, surface waters) were then merged into a single table spanning the three distinct environments.

Since the primer pair used to PCR amplify a portion of the *hsp65* gene (Tb11-Tb12) is not specific to members of the genus *Mycobacterium*, it was necessary to remove all non-mycobacterial reads. The combined ASV representative sequence file (repset) generated above was then filtered using the updated NTM database as the reference database using blastn (Altshul et al, 1990)(command: makeblastdb -in < NTM DB >)(see below for details) at a minimum match percent identity of 90% or greater. The combined ASV repset file was then filtered using *seqtk* (https://github.com/lh3/seqtk) to retain only ASVs that met the 90% or greater match threshold to the reference database.

Samples with fewer than 100 mycobacterial *hsp65* reads were removed from the dataset, resulting in a total of 822 samples total across the three environments (household plumbing biofilm N= 507, soil N= 140, surface waters N= 175). This table was used in all downstream NTM analyses. Across these 822 samples, we detected a total of 7,854 mycobacterial ASVs.

### Updated hsp65 marker gene database and phylogenetic tree

The NTM phylogenetic tree was built using an updated version of the Dai et al. 65-kD heat shock protein marker gene database (Dai et al. 2011). Briefly, the database was aligned and trimmed using MAFFT (version v7.505 (2022/Apr/10)(Katoh et al. 2002) and trimal (v1.4.rev15 build[2013-12-17])(Capella-Gutiérrez, Silla-Martínez, and Gabaldón 2009), respectively. The aligned database was then used to train a Hidden Markov Model using HMMR (HMMER 3.3.2 (Nov 2020))((Wheeler and Eddy 2013)). The HMM was then used to search (hmmsearch) a curated file of mycobacterial whole genome sequences extracted from the Genome Taxonomy Database (GTDB)(Parks et al. 2022). To remove any sequence noise and mismatches from the output file, a phylogenetic tree was built using RAxML (version 8.2.12), and divergent sequences were removed from the database. The final database resulted in 189 mycobacterial heat shock protein sequences, adding an additional 32 NTM species or strains to the 2011 Dai et al database. The updated reference database can be found at https://github.com/gebertm/hsp65_Tb11-Tb12_db.

### ASV Clustering

All 7,854 mycobacterial ASVs were clustered to combine ASVs that were closely related based on phylogenetic analyses. This was done as we observed a high diversity of individual ASVs and because most ASVs were only found in a relatively small number of samples across all three datasets (mean of 6.4 samples per ASV). To conduct the phylogenetic clustering, we built a phylogenetic tree, including both the representative sequence file and updated *hsp65* marker gene database, to assign taxonomy based on proximity to a known reference database sequence, using *Nocardia farcinica* (DSM43665) as the outgroup. Briefly, sequences were aligned using MAFFT (Katoh et al. 2002) (v7.505 (2022/Apr/10)), and aligned sequences were then trimmed using trimAl (v1.4.rev15 build[2013-12-17]). The maximum likelihood tree was built using RAxML (version 8.2.12, 2018) with the following command raxmlhpc -f a -m GTRGAMMA -p 12345 -x 12345 -# 100 -T 30 --print-identical-sequences.

To cluster ASVs, we used the TreeCluster package (Balaban et al. 2019) (version 1.0.4) with the distance threshold t = 0.20, meaning 0.20 is the maximum pairwise distance between leaves in a given cluster. The optimal threshold was set based on the clustering of the *Mycobacterium tuberculosis* complex reference sequences into one unique cluster, without including additional NTM taxa. TreeCluster resulted in 404 unique clusters, encompassing the 7,854 ASVs. For a cluster to be present in a sample, it was required to have a total of >10 reads across the combined dataset. 62 ASVs did not fall within a defined cluster (62/7854, or 0.79%), designated as *-1* in the final cluster table (Supplementary Table 2), and were excluded from the downstream analysis. Taxonomic names were assigned at the cluster level, when possible (i.e., clusters that included named reference strains were assigned the name of that reference strain). Only 74/404 clusters (18%) contained one or more reference sequences, meaning most clusters (82%) could not be confidently assigned to the species level based on their phylogenetic similarity to a named species.

The top 50 clusters found in each environment, which encompassed >99%, 66%, and 95% of the total reads in household plumbing biofilms, soil, and surface waters, respectively were selected for downstream analyses. Since the NTM diversity in the soil samples was considerably higher compared with the other two environments, a smaller percentage of the total NTM reads from the soil samples were captured in the top 50 clusters.

### Statistical Analysis

To visualize the variation in community structure of all NTM species, we ran a principal components analysis (PCoA) on the cleaned cluster table using the Phyloseq package in R (McMurdie and Holmes 2013). Briefly, we converted the cluster table from read counts to a presence/absence matrix, requiring a minimum of 10 reads per cluster for it to be considered ‘present’ in the data. The dissimilarity matrix was calculated using pairwise Jaccard distances.

To test whether any of the measured environmental variables collected at each USGS site predicted the NTM community composition in the surface water samples, we ran a Mantel test on samples using the R vegan package (version 2.6-8) (Oksanen et al. 2013). Briefly, for the measured environmental variables, an additional dissimilarity matrix was calculated using *vegdist* in the R vegan package (version 2.6-8) (Oksanen et al. 2013) using pairwise Euclidean distance. A Mantel test was then run between the taxa dissimilarity matrix and the metadata dissimilarity matrix to determine if any of the measured environmental variables predicted variation in NTM community composition across samples.

## Results

### Relative abundance of genus Mycobacterium across the three environments

We used the 16S rRNA gene sequence data to determine the relative abundance of the genus *Mycobacterium* across the 1,154 samples from the three environments (Figure 2). Members of the genus *Mycobacterium* were nearly ubiquitous across the three environments, detected in 100% of the 146 soils, 76% of the 373 surface water samples, and 94% of the 638 household plumbing biofilms. However, the relative abundances of mycobacteria varied depending on the environment. In household plumbing biofilms, mycobacterial abundances ranged from no mycobacteria detected (40 samples) to mycobacteria representing >95% of the bacterial communities (4 samples). The median relative abundance of the genus *Mycobacterium* in the plumbing biofilms (3.7%) was appreciably higher than that observed in soil (median relative abundance = 0.37%) and surface waters (median relative abundance = 0.015%). Only 1.3% of surface water samples (5/373) and 11% of soil samples (16/143) had mycobacterial abundances over 1%, while 60% of household plumbing biofilm samples had mycobacterial abundances greater than 1% (384/638) (Figure 2).

**Figure 2:**
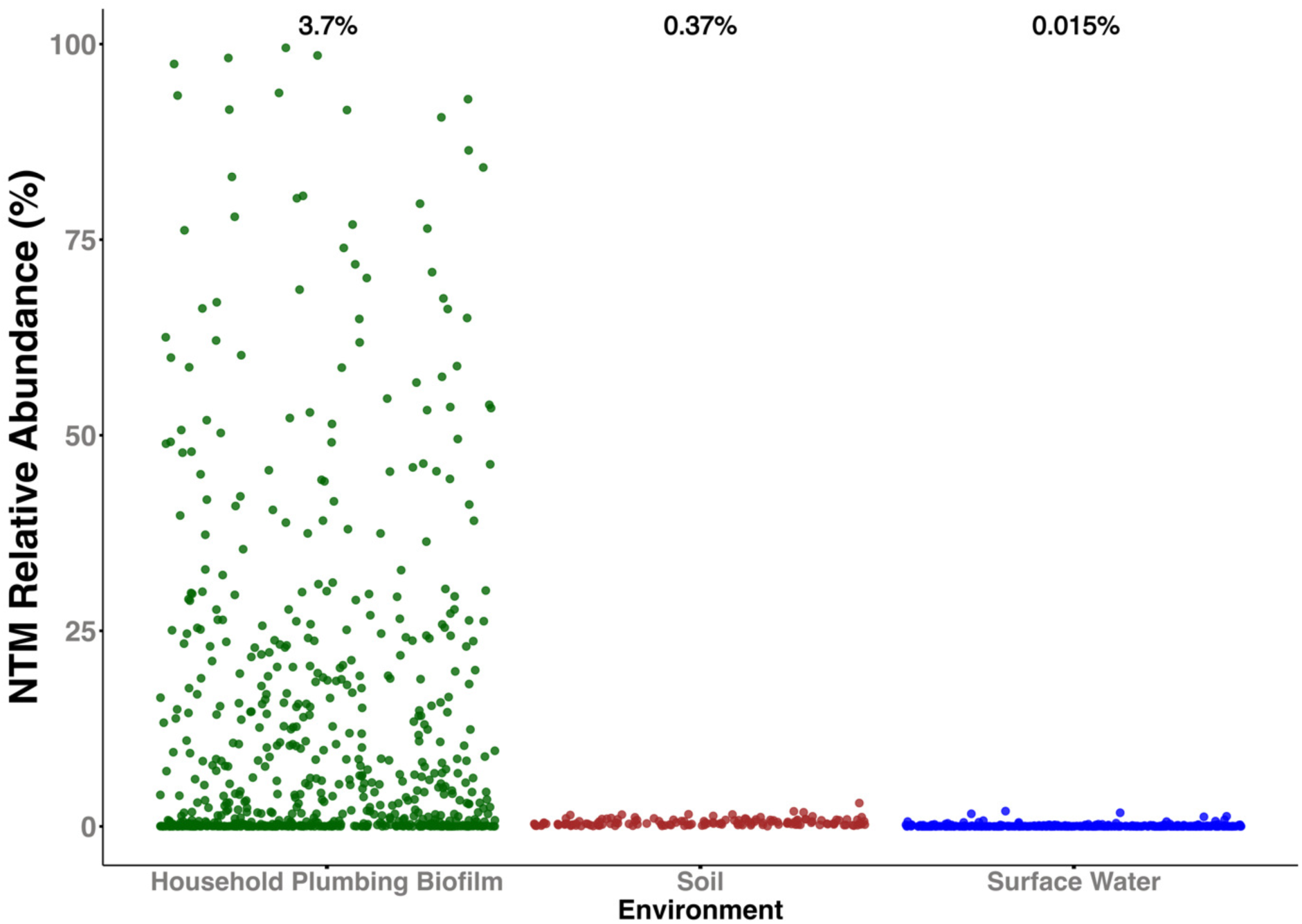
Relative abundances of the genus *Mycobacterium* across three environments (household plumbing biofilm (n=638), soil (n=143), and surface water (n=373)). Relative abundances represent the proportional abundances of taxa assigned to the genus *Mycobacterium* compared to all other bacteria and archaea detected in each sample from 16S rRNA gene sequencing effort. Median relative abundances of the genus in each environment are noted above each scatterplot.

### Mycobacterial species-level survey across the three environments

While 16S rRNA marker gene sequencing allowed us to compare the relative abundance of the genus *Mycobacterium* across environment types, species-level information obtained through mycobacterial-specific marker gene sequencing of the *hsp65* gene, is crucial for understanding the diversity of mycobacterial taxa in each environment, especially those of clinical significance. To better summarize the broad-scale diversity of the NTM in these environments, we phylogenetically grouped mycobacterial ASVs into “clusters” based on phylogenetic relatedness. Most NTM clusters found in the household plumbing biofilms clustered with known reference sequences (>80% sequence similarity to an isolate found in the reference database), including many of known clinical significance (for example *Mycobacterium avium* and *Mycobacterium chelonae*). In contrast, many NTM clusters identified in soil and surface waters could not be identified below the genus level of taxonomic resolution as they did not include any named species in our phylogenetic analyses. When we focused on the top 50 NTM clusters detected in each of the three environments (150 in total), we found that in household plumbing biofilms, 24 clusters were found to contain a reference sequence (48%), but only 24% of clusters in soil and 16% of clusters in surface waters shared similarity with a reference sequence. All three environments harbored a broad diversity of NTM, spanning the mycobacterial phylogenetic tree (Figure 3), however, the amount of novel taxonomic diversity observed in natural environments was higher than what was observed in household plumbing biofilms.

**Figure 3:**
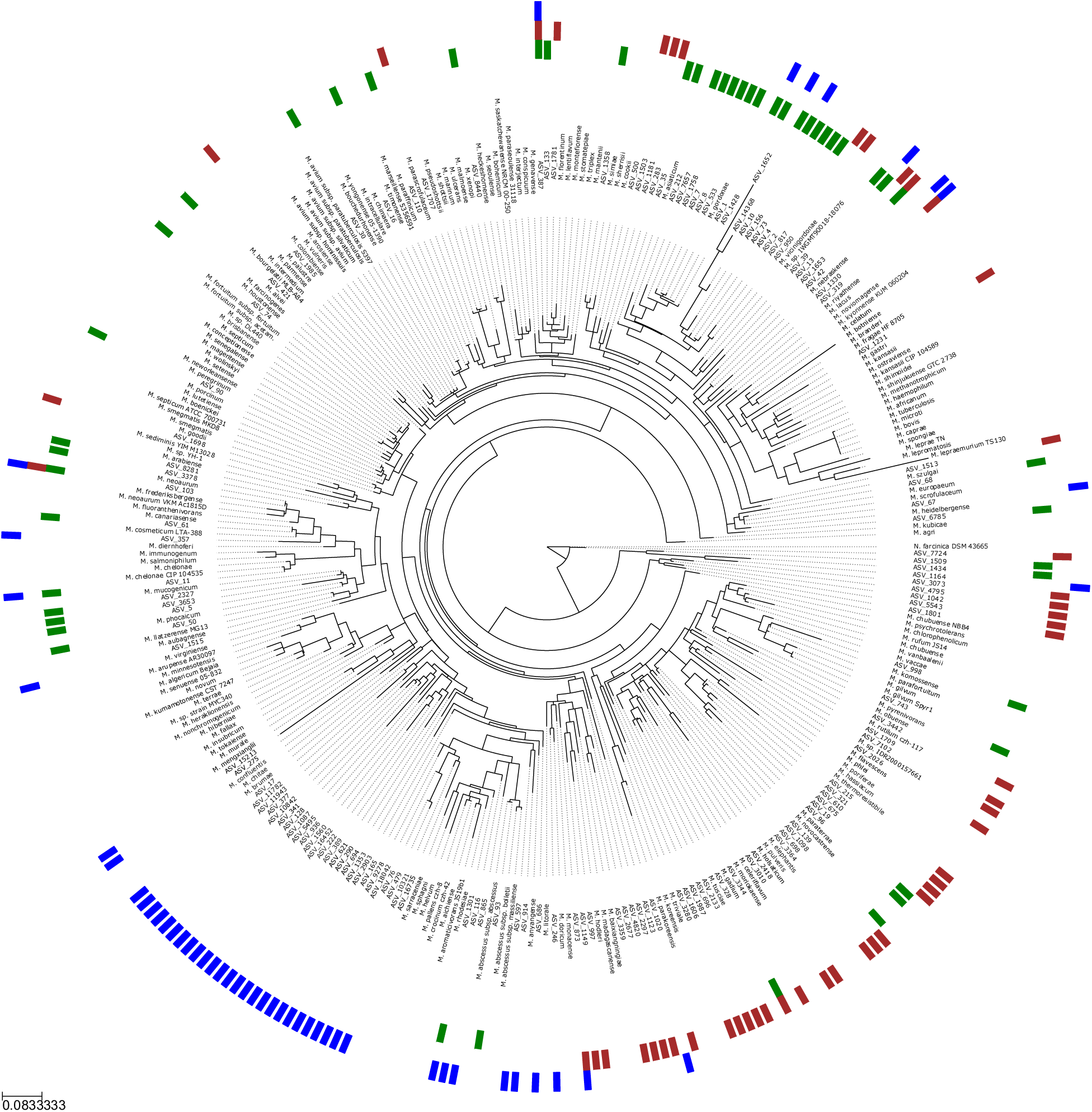
Maximum likelihood tree, using the mycobacterial *hsp65* gene, showing 189 reference strains, plus one representative ASV from each of the top 50 NTM clusters detected in each environment (135 unique ASVs total). Colors of outside ring represent the environment in which the NTM cluster was detected (household plumbing biofilm = green, soil = brown, surface waters = blue). Phylogenetic tree was rooted using the *hsp65* sequence from *Nocardia farcinica* DSM43665.

### NTM Cluster Occupancy

Although NTM were detected in nearly every sample, regardless of the environment in question, the composition of the mycobacterial communities varied across environment types (Figure 4). The most abundant NTM taxa found in the household plumbing biofilm samples were *M. gordonae*, followed closely by *M. mucogenicum/phocaicum*, while the most abundant taxa found in soil were *M. madagascariense* and *M. holsaticum*. The most abundant NTMs in the surface water dataset could not be classified to species as they were not closely related to any named NTM species in our reference database (Figure 4C). The variation in NTM community composition across the household plumbing biofilm samples was far greater than the variation detected across either the soil or surface water samples (Figure 4). Additionally, the composition of the NTM communities found in surface waters and soils were generally more similar to each other than to those found in the household plumbing biofilm samples (Figure 4), highlighting that each of the three broad environment types harbor distinct NTM communities.

**Figure 4:**
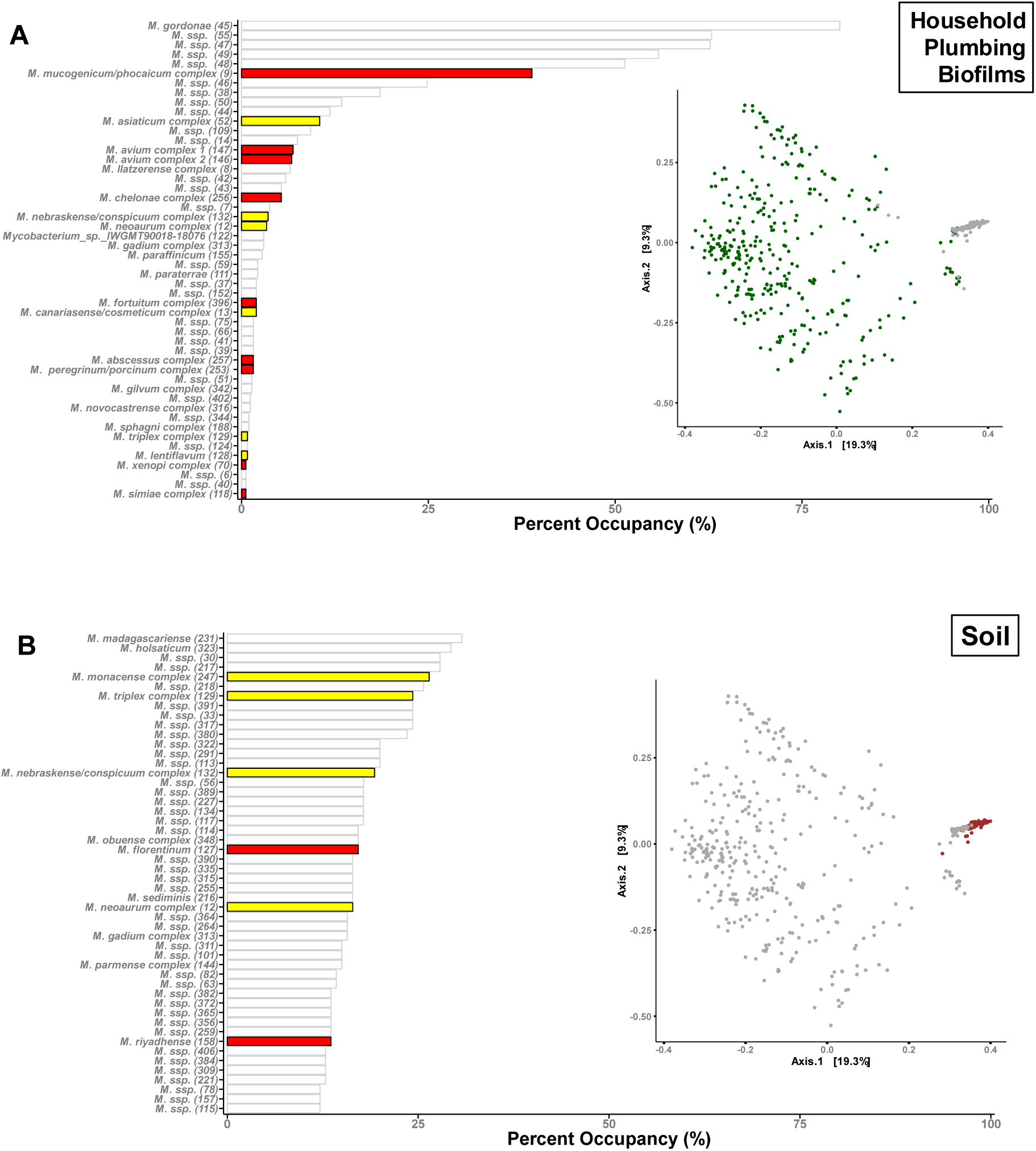
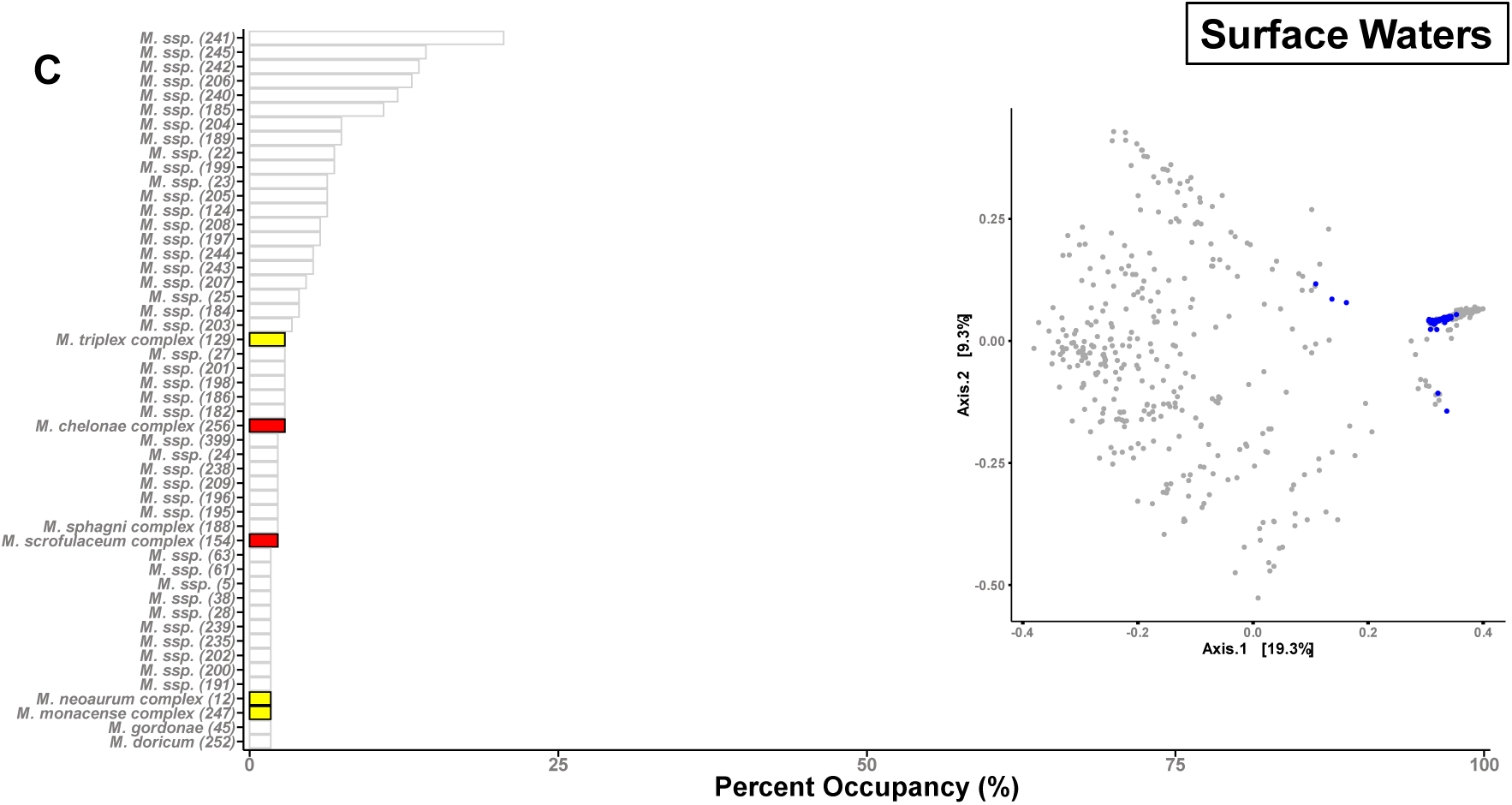
Bar plots showing the percent of samples (percent occupancy, i.e. percent of samples from each environment in which a given taxon was detected). Panel A: household plumbing, Panel B: global soils, Panel C: surface waters. Bars shaded in red are NTM taxa inferred to have high clinical significance, while bars shaded in yellow have low, or limited, clinical significance. Bars outlined in gray have no reported clinical significance. Taxa indicated by 8*M. ssp.*9 represent lineages that do not include any named mycobacterial isolates. On the right of each bar plot are shown principal coordinate analysis (PCoA) colored by environment type (household plumbing biofilms in green, soils in brown, surface waters in blue), showing variation in overall NTM community composition across the three environments. We note that that three plots are identical except for the highlighting of specific sample types.

### Relative abundance and ubiquity of clinically relevant NTM in the environment

The number of samples that contained clinically relevant NTM varied between environment types, with most clinically relevant taxa detected in the household plumbing biofilm samples (*M. avium* complex, *M. mucogenicum*, *M. abscessus*), while the top 50 taxa detected in soils only contained a handful of named strains, two of which have limited clinical significance (*M. florentinum* and *M. riyadhense*, detected in 17% and 14% of samples, respectively). The surface water samples contained only two named clinically relevant NTM, *M. chelonae* complex and the *M. scrofulaceum* complex, which were detected in 2.9% and 2.3% of samples, respectively.

We chose to focus on six clinically relevant nontuberculous mycobacterial taxa that are frequently detected in clinical samples from human patients (*M. mucogenicum*/*M. phocaicum*, *M. avium* complex, *M. chelonae* complex, *M. abscessus* complex, and *M. fortuitum* complex, (Daley et al. 2020)) and how their relative abundances and occurrence patterns varied across the three environments (Figure 4). All six clinically relevant NTM clusters were detected in household plumbing biofilms. Four of the 6 taxa (*M. mucogenicum/phocaicum*, *M. avium* complex 1, *M. chelonae* complex, and *M. abscessus* complex) were the most ubiquitous in household plumbing biofilms (Figure 5). For example, *M. mucogenicum/phocaicum* was found in ∼39% of samples from household plumbing biofilms, with a mean relative abundance of 16% of mycobacterial *hsp65* reads, compared to less than 1% in the natural environments. The *M. avium* complex 1 was not detected in either soil or surface water samples but was found in nearly 7% of the household plumbing biofilm samples, with a mean relative abundance of 1.7% (Figure 5). The *M. avium* complex 2 and the *M. fortuitum* complex, although not detected in surface waters, were detected in a greater number of soil samples compared to household plumbing biofilms (10% vs. 6.9% and 12.1% vs. 2%, respectively). Although the *M. avium* complex 2 was detected across more soil samples, it had a higher mean relative abundance in household plumbing biofilms (1.4%), compared with soil (0.29%).

**Figure 5:**
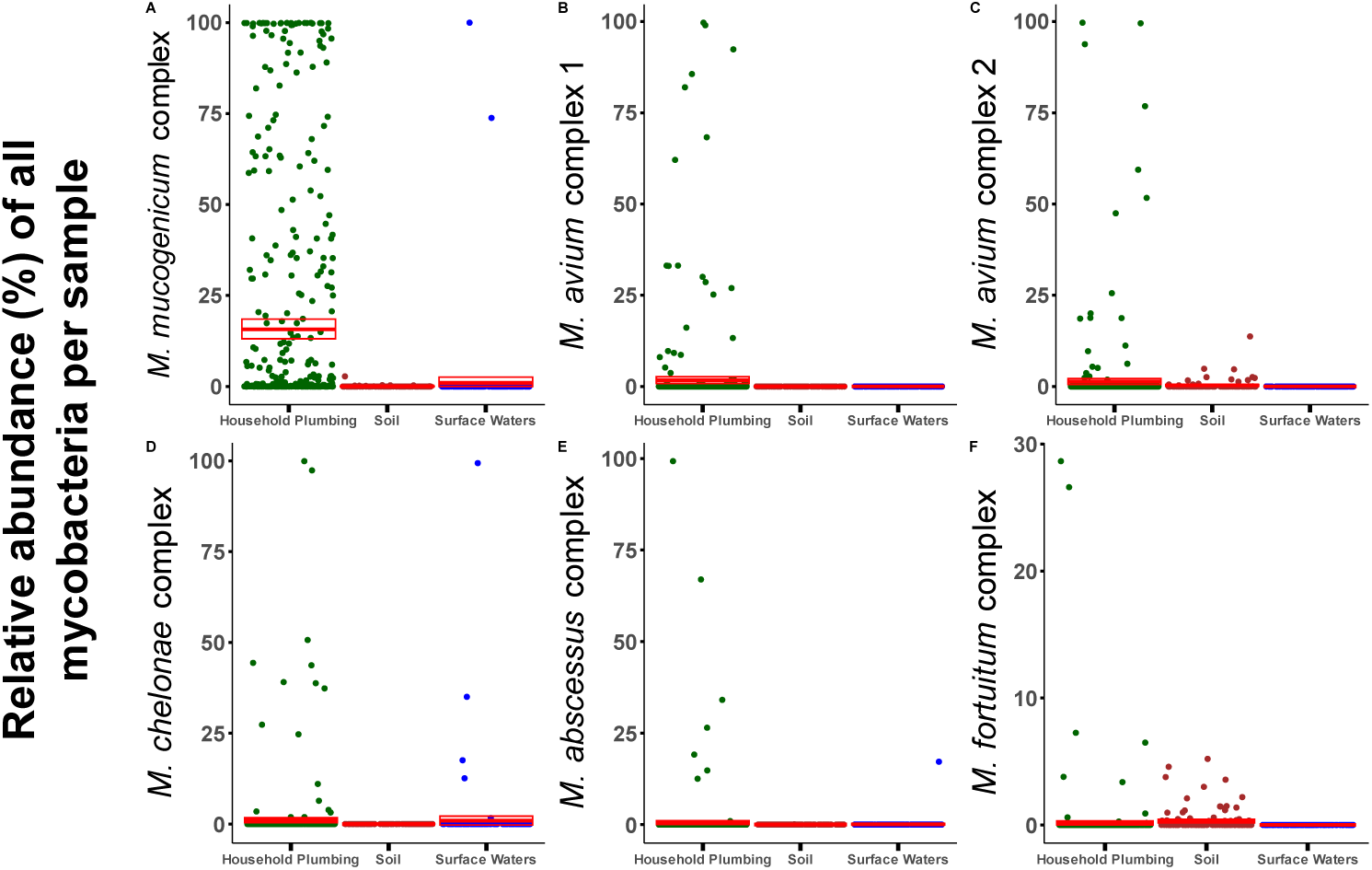
Scatterplots showing the relative abundance of six clinically relevant NTM groups across the three environments: A) *M. mucogenicum* complex, B) *M. avium* complex 1, C) *M. avium* complex 2, D) *M. chelonae* complex, E) *M. abscessus* complex, F) *M. fortuitum* complex. Mean relative abundances and 95% confidence intervals are shown in red. Relative abundances were calculated by dividing the number of *hsp65* reads assigned to each group in a sample by the total number of mycobacterial *hsp65* reads from that sample.

## Discussion and Conclusions

Nontuberculous mycobacteria pulmonary disease is on the rise in the United States and around the world (Dahl et al. 2022; Bents et al. 2024). Although many environments have been implicated in the direct transmission of disease, the ecology of NTM across environments remains poorly understood, and studies showing how NTM taxa, including known pulmonary pathogens, vary between environment types remain limited. Here we characterized the variation in the amounts and types of NTM, including those of clinical significance, across a broad range of environments frequently implicated as sources of NTM exposure, including both natural environments (soils, surface waters) and a built environment (household premise plumbing).

While we detected members of the genus *Mycobacterium* in nearly all samples from all three environments, the relative abundance and types of NTM differed appreciably across environments. The median relative abundance of the genus *Mycobacterium* in both natural environments was low compared to that of household plumbing. We also found that the three environments harbored diverse and distinct NTM communities (Figure 3), but most of the NTM taxa identified, particularly in the natural environments, were of limited to no clinical significance. Of note, clinically significant NTM (including *M. mucogenicum* and *M. abscessus*) were more frequently detected and more abundant in household plumbing biofilms compared to the natural environments surveyed (Figure 4). These results suggest that household plumbing biofilms, not soils or surface waters, likely pose a higher risk of NTM pathogen exposure, and subsequent infection. Although clinically relevant NTM were more frequently detected in household plumbing samples, we note that detection of pathogens does not necessarily equate to a higher risk of infection from household water sources as association between the number of pathogenic NTM cells and disease risk remains uncertain (Falkinham 2016). Likewise, differences in exposure routes, duration, and frequencies likely influence infection risk. However, our results do suggest that patients at risk of NTM infections are more likely to acquire infections from premise plumbing sources than from soils or surface waters, emphasizing the potential risks associated with exposures to certain environments.

Of note, the *M. scrofulaceum* complex, which encompasses the clinically relevant species *M. scrofulaceum*, *M. interjectum*, and *M. malmoense*, appears rarely in the household plumbing biofilm samples (occupancy = 0.39%) but appears more frequently in both natural environments (occupancy = 10% in soil, and 2.3% in surface waters), reflecting the known susceptibility of this NTM group to the chlorination practices commonly used to treat household water supplies in the US, practices that may be associated with a decrease in *M. scrofulaceum* cases in the US (Falkinham 2003).

Together our study highlights that, while many environments harbor NTM, they do not all harbor the same amounts and types of NTM, including clinically relevant taxa which are particularly ubiquitous and abundant in premise plumbing biofilms compared to soils and surface waters. Overall, a better understanding of the ecology of NTM in source environments, including the distributions of clinically relevant NTM taxa, should help inform epidemiological investigations of NTM disease outbreaks and guide recommendations provided to disease susceptible individuals to minimize NTM exposures. Future research should focus on the factors that contribute to NTM selection in the built environment – a likely important route of exposure for human disease. Understanding where certain NTM, especially potential pulmonary pathogens, are coming from, will play a key role in potentially preventing their accumulation in premise plumbing biofilms.

## Acknowledgements

We thank Caihong Vanderburgh for her assistance in the laboratory with sample processing, Holly Roth for help with remote access to the USGS databases, Dr. Andrew Blakney for providing feedback on the manuscript, as well as everyone at USGS collection sites across the United States who contributed filters to this study. Additionally, we thank Dr. Conor Meehan for his assistance and guidance with the updated *hsp65* database.

